# Towards an ontology-based recommender system for relevant bioinformatics workflows

**DOI:** 10.1101/082776

**Authors:** Ahmed Halioui, Petko Valtchev, Abdoulaye Baniré Diallo

## Abstract

**Background:** With the large and diverse type of biological data, bioinformatic solutions are being more complex and computationally intensive. New specialized data skills need to be acquired by researchers in order to follow this development. Workflow Management Systems rise as an efficient way to automate tasks through abstract models in order to assist users during their problem solving tasks. However, current solutions could have several problems in reusing the developed models for given tasks. The large amount of heterogenous data and the lack of knowledge in using bioinformatics tools could mislead the users during their analyses. To tackle this issue, we propose an ontology-based workflow-mining framework generating semantic models of bioinformatic best practices in order to assist scientists. To this end, concrete workflows are extracted from scientific articles and then mined using a rich domain ontology.

**Results:** In this study, we explore the specific topics of phylogenetic analyses. We annotated more than 300 recent articles using different ontological concepts and relations. Relative supports (frequencies) of discovered workflow components in texts show interesting results of relevant resources currently used in the different phylogenetic analysis steps. Mining concrete workflows from texts lead us to discover abstract but relevant patterns of the best combinations of tools, parameters and input data for specific phylogenetic problems.

**Conclusions:** Extracted patterns would make workflows more intuitive and easy to be reused in similar situations. This could provide a stepping-stone into the identification of best practices and pave the road to a recommender system.

## 1 Background

The advances in biotechnology have revolutionized bioinformatics into a data-intensive discipline. The rapid increasing amounts of data available from new large scale analyses have made data processing without automated pipelines (*a.k.a.* workflows) difficult. This situation is mainly due to the complexity of integrating diverse data formats and databanks, interfacing numerous programs with various parameters, as well as assessing software and data. Current bioinformatic platforms offer a variety of computational solutions (by high-performance and cloud computing) with different levels of computational abstractions [1]. Two types of models are employed by workflow systems: concrete ones, in which the tasks are bound to specific resources, and abstract ones, presenting an abstract view of the resources used in the tasks. Nowadays, abstract bioinformatic workflows offer only service-oriented platforms such as Pegasys [2], Taverna [3], Galaxy [4] or Armadillo [5]. However, none of them provide recommendations to guide users and suggest steps while taking into account the specific nature of data (complete vs partial genomes, viral proteins, plant DNA, type of the evolutionary model, *etc.*) and relevant analyses (phylogenetic analyses, function prediction, homology studies, genome wide association studies, etc.).

In phylogenetic tree reconstruction, the evolutionary history of a group of organisms could be inferred with different types of data including nucleic acid (DNA or RNA) or protein sequences. Depending on the used tools, different types of evolutionary events could be included in the study such as point mutations, duplications, insertions and deletions, genome rearrangements, etc. [6]. Such a study requires several difficult steps from the acquisition of the raw data. Phylogenetic analysis constitutes a typical bioinformatic workflows in life sciences. This is witnessed by the large number of publications on the topic. This resource could be exploited to enrich problem solving solutions. The difficulties in the process of construction of a phylogenetic workflow from these publications are illustrated in the following example.

### Example 1

A biologist *U*_1_ downloads a set of 9 hemagglutinin sequences of different Influenza viruses from the NCBI database. To carry out the analysis of those sequences, *U*_1_ chooses the Armadillo [5] platform[^1^]. Several algorithmic options are displayed: i.e. software parameters (*e.g.* the number of bootstraps, mutation rate, *etc.*) and data constraints (*e.g.* input/output to programs). *U*_1_’s prior *abstract* experience doesn’t cover all the available methods and tools, therefore the backup solution is to adapt a workflow from the literature (see Fig. 1). However, none of the workflows drawn from scientific publications could exactly match his needs. These workflows may not properly execute since a part of the analysis requires several *specific* software versions and dependencies.

**Figure 1.**
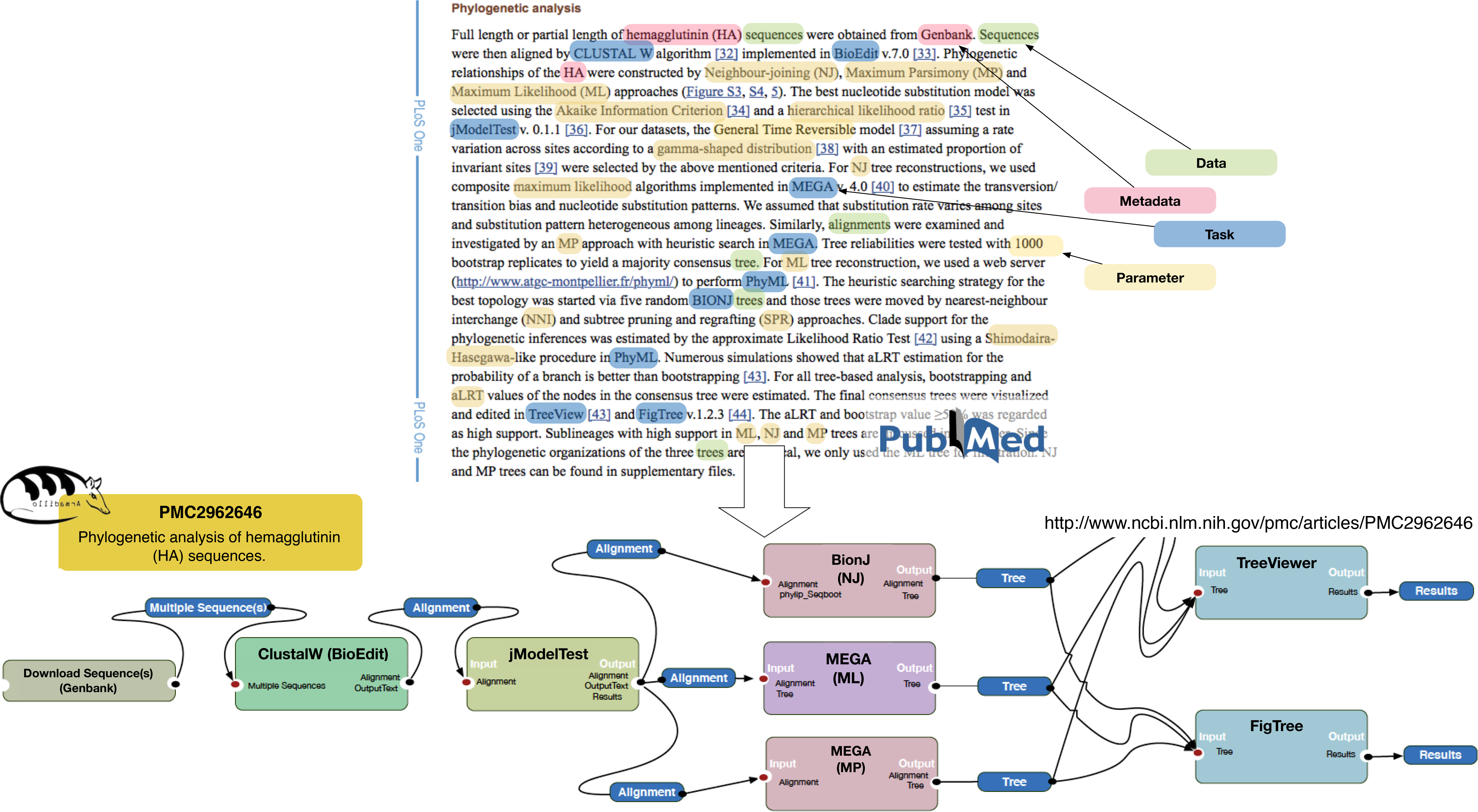
A Phylogenetic analysis example drawn from the publication [47] using the Armadillo [5] platform. From the highlighted softwares, data, descriptive metadata and parameters in texts, an expert drew a phylogenetic workflow. Task programs are represented with coloured rectangles and data flow with labeled links (input/output parameters).

The above example shows several problems which could occur while constructing bioinformatic workflows. Even simple routine projects can involve a number of different softwares requiring specific but also abstract problem solving knowledge. Even though software experimental conditions are related to several choices, the amount of data and the lack of knowledge of tools could also deteriorate the results of studies [7]. This is where users’ skills and experience do make the difference. Due to this kind of lack of knowledge related to experimental conditions and analysis tools choices, researchers could end up confused or unable to explain their choices [7]. To tackle these issues, we propose to explore more multi-level abstract workflows in order to assist users during their tasks. We propose an ontology-based approach towards extracting and formalizing of bioinformatics practices, specifically phylogenetic data skills, from the specialized literature.

### Example 2

Suppose that a bioinformatician *U*_2_ chooses to study the same protein sequences of the hemagglutinin. He runs *ClustalX* to align sequences and then chooses the *JTT* evolutionary model during the tree reconstruction. If we want to recommend a concrete phylogenetic program to *U*_2_ such as the program *MEGA,* we propose to calculate the frequency of the sequence *s*_0_ : 〈 *ClustalX, JTT, MEGA* 〉 in the literature of thousands of tools and versions. However, such sequence is very rare due the vast varieties of available tools for the specific task: phylogenetic inference. Thus, recommending a category of tools such as any “maximum parsimony” or “maximum likelihood” approach is a more relevant recommendation. An abstract pattern such *p*_0_ : 〈 *Local Alignment, ProteinModelSelection, MaximumLikelihood* 〉 is more frequent than the concrete pattern *s*_0_. Here, we use ontologies to represent the different categories (concepts) of tools and resource annotations at different levels of abstractions. The constraints on combining tools and parameter values are expressed with properties (relations) between classes. The ontological representations are effectively exploited by two key processes in our approach: (1) the extraction of workflow terms and relations from texts (*concrete workflows*), and (2) the extraction of generalized workflow patterns (*abstract workflows*). Here, we distinguish concrete workflows made of individuals of our ontology and links between them, from abstract ones comprising of concepts and properties at various levels of abstraction. Problem-solving patterns mined from concrete examples, could be the foundation of recommending best practices within a workflow-based environment.

Current bioinformatics Workflow Management Systems (WMS) have emerged as global *intelligent* infrastructures by integrating new technologies to maintain a certain level of reusability through grid and cloud computing [1, 8]. Yet, we consider intelligence, a system to help compose workflows, which we define here as guided assistance based on the context of the bioinformatics problem-solving. Our level of understanding should better solve problems and make decisions via Knowledge Management (KM) discipline. KM is the subject that provides strategy, process and technology to share information and expertise among users. For instance, Semantic Web (SW) technologies [9], as a special case of KM, allow automatic discovery and execution of web services that can handle the workflow tasks. Similarly, ontologies have substantially improved knowledge integration, querying, and sharing in the bioinformatics domain [10]. Bioinformatic workflow systems could benefit from the SW efforts to develop strategies to capture knowledge about the available bioinformatics tools, services and algorithms. Proteus [11] is one of the first projects to integrate an ontology-based design in bioinformatic workflows. It exploits a *DAML+OIL* ontology for the data mining domain describing resources and processes of knowledge discovery in databases. However, formalizing bioinformatics solutions in a hierarchy of applications is not enough if there is no reasoning engine to make benefits of the presented semantics to assist and guide users. Proteus provides semantics about resources but doesn’t recommend the best categories for a specific task. In fact, requirements and technologies for bioinformatic applications should allow the WMS to specify complex problem solving applications. Current solutions for bioinformatics recommendation can be grouped into three categories: (1) expert recommendation, (2) semantic-based recommendation and (3) pattern-based recommendation.

### Experts recommendation

In the literature of bioinformatic best practises, few works report on experiences and/or provide suggestions for the current WMS. In this vein, Spjuth et al. [1] review the most advanced projects defining general guidelines for the methodological choices at various stages of a bioinformatics pipeline. The authors recommend dedicated practices in maintaining reproducibility and standardization in local and distributed environments. Anisimova et al. [6] criticize the outdated methodology and provides practical guidance on issues and constraints faced by each of the phylogenetic analysis steps. Garijo et al. [12] propose a set of high-level abstract motifs (patterns) as a view of activities undertaken within workflows. They performed their manual analysis over a set of real-world scientific workflows from Taverna and Wings systems.

### Semantic-based recommendation

Lee et al. [13] describes a SW solution assuming a Grid computing environment. The proposed Bio-STEER system offers SW services in OWL-S. A small-size ontology is employed in the representation of both abstract workflows (for Grid resources) and concrete ones (to bind tasks to specific resources). Ison et al. [14] introduce the EDAM ontology for bioinformatic operations, types of data, identifiers, topics and formats. EDAM supports semantic annotations for web services, databases, interactive tools and libraries. The ontology has been used to find, describe, compare and select tools into workflows. EDAM annotations have been implemented in a set of frameworks such as EMBOSS ^[2]^ and eSysbio ^[3]^. However, these systems don’t exploit the entire richness of EDAM semantics. For example, eSysbio uses “EDAM Data and Format” to decide how to handle data by an adequate visualization and search. However this navigation, as static as it is, doesn’t take the advantage of the relations between concepts. Navigating and searching the right tools in categories of tools are not enough to guide users to complete their tasks. Formulating queries could be a very difficult task if users don’t know what is the best category for the next step. Digiampietri et al. [15] propose the SHOP2 specification which comprises three sections: a domain definition (defdomain), a problem definition (defproblem) and a problem resolver (find-plans) where the latter is being in charge of finding the plans to solve the targeted problem. The respective SHOP2 planner supports generalization/specialization hierarchies of operators (tasks) and handles complex objects: structures created from basic objects by composition. It uses ontologies to semantically support workflow construction for a better selection of appropriate tasks and services.

### Pattern-based recommendation

Pattern-based recommenders use sequential pattern mining algorithms [16] in order to discover recurring software and data usage patterns. Duck et al. [17] propose to use Natural Language Processing (NLP) tools to extract software and database terms from scientific texts. The process uses the bioNerDS terminology ^[4]^ in order to categorize terms in texts. In the pattern mining step, a straightforward mining algorithm discovers frequently occurring resource pairs. Each resource is paired with the one that immediately follows it in a text while accounting for the direction within the ensuing pairs. However, the order of the terms in the text is not always reliable. Without a context-aware extraction approach, the occurrence of such a pair could mean nothing but simple mentions. Soomro et al. [18] propose a pattern-based recommender system for neuroimaging workflows. While the described implementation is domain-specific, the architecture could easily be reused in bioinformatics workflows. The workflows are first converted into graphs, their components are then generalized and the result is fed into a pattern miner. Once matching patterns are found, they are specialized and the partial workflow (the workflow to recommend) is semantically analyzed and enriched. Authors don’t provide details about their pattern mining algorithm but show interesting results in terms of Mean Reciprocal Rank (MRR). Other measures to calculate workflow similarities are proposed in [19–21]. In the same scope of graph-based workflow discovery, Goderis et al. [22] attempt to build a gold standard for workflow ranking based on graph sub-isomorphism matching. However, their method have been only tested on 89 different workflows.

Lord et al. [23] describe a classification of bioinformatics workflows that could be used to suggest classes of tools while creating a workflow. Their proposed approach, based on clustering methods, considers program terms but also some structural workflow informations (pairs of tasks) and input-output parameters in the encoding phase. However, taking into account these informations doesn’t improve the quality of workflow classification due the spareness of data encodings (using matrices). In a different vein, methods for mining patterns and associations on top of an ontology have been proposed in [24]. As an example, the GSP (Generalized Sequential Patterns) algorithm [25] mines frequent patterns from sequences of user transactions. It traverses the pattern space with an Apriori-like level-wise discipline [26] and by exploring the monotony of itemset frequencies over the taxonomy. Besides, using an ontology to enrich patterns of Web usage has been researched on for at least a decade [27, 28]. However, these latter only process the class hierarchy.

Here, we follow the methodology described in [29] where the input data are sequences of clicks further translated into IDs of content objects with their semantic links. Our own approach could be seen as an extension thereof where, beside workflow sequences extracted from texts, it also comprises labeled links between items, thus, forming a layered 2-labeled DAG (Direct Acyclic Graph). Our proposed workflow language is both richer (various sorts of interdependencies) and less constrained (lack of a total order on workflow elements) hence costlier to process. Thus, we put the emphasis on pattern-to-data matching and on parsimony of pattern space traversal. In our case, on top of the workflow DAGs, an ontology provides two taxonomies, one for concepts (a.k.a. generalized items) and one for relations. Hence, our patterns are made of concepts with inter-concept relations which may belong to different generality levels in the ontology.

The remainder of the paper is organized as follows: section 2, we define the pipeline of our approach. We also describe the methods for concrete workflow extraction from texts and for generalized pattern mining. Section 3 presents the experimental evaluation and results of the extracted patterns. We discuss the quality of our approach in section 4 and conclude in the last section.

### 2 Methods

The proposed solution consists of reconstructing a set of workflows from the scientific literature and then mining higher-level of abstractions (generalized workflow patterns) from these. A specific domain ontology is used to extract concrete workflows from texts. The included concepts and properties are further used as building blocks in the pattern generation process. The ontology is exploited for two purposes: (1) it is used as schema to create a terminology to annotate workflows in texts and (2) as a knowledge base to mine generalized patterns from the extracted ones.

#### 2.1 Formalisms for enriched workflow representations

##### 2.1.1 Workflow representation

In general, a workflow consists of a set of tasks (also called activities) combined in a *control-flow* and a *data-flow* whereby the data-flow describes interactions (relationships) between tasks and data items (task inputs/outputs corresponding to data types). Given a universe of task items and data *O* and links 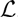, a workflow record is a vertex- and edge-labeled DAG. However, by following the steps of the overall analytical process -and performing a topological sort, if neededit can be split into layers of independent vertices (no intra-layer edges). This allows a workflow representation as: (1) a sequence of object sets (item sets), plus (2) a set of cross-itemset links.

A data language ∆_*W*_ defines workflows as pairs *w* = (*ζ,* Θ) where 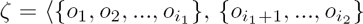, 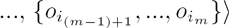 is the sequence of object *transactions* (steps). In turn, 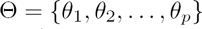 is the set of triples where *θ*_*k*_ = (*i*_1_, *l*, *i*_2_) represents a link of type *l* between the object at position *i*_1_ in 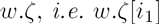 and at *w.ζ*[*i*_2_] where *i*_1_ < *i*_2_. Fig. 2 shows an example of workflow representation in ∆_*W*_.

**Figure 2.**
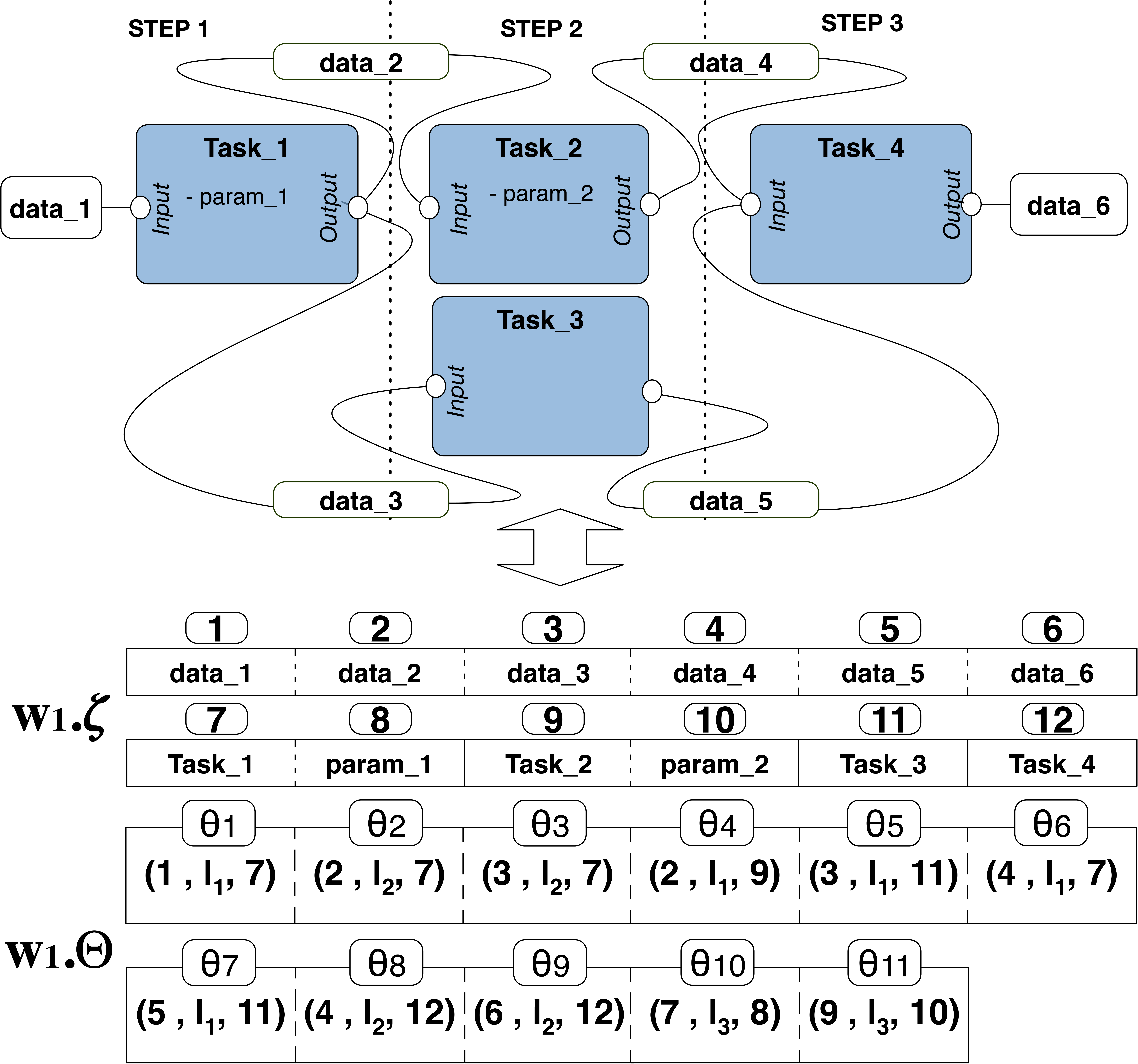
An example of workflow representation. Workflow items: *i.e.* tasks, data and parameters are encoded in the sequential set *w*_1_.*ζ* while links between them are encoded in the triple set *w*_1_.*Θ*. Each triple represents a link between item positions. Here, the links *l*_1_, *l*_2_ and *l*_3_ represent respectively, the *isInput, isOutput* and *hasParameter* relations

##### 2.1.2 Ontology representation

Workflow components are encoded in a specific domain ontology Ω. Formally, an ontology is defined as a six-tuple 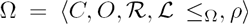 where *O* is the set of all object items, *i.e.* individual tasks and data types, 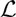 the set of all links between objects, *C* the set of all concepts, and 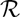 the set of all relations between concepts. Moreover, concepts and relations are organized in taxonomies *w.r.t.* the generality order *(subClassOf* relationship in RDF) of the ontology 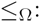*i.e.*: 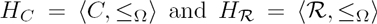, the hierarchical order in the ontology. 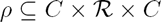 is a ternary relation whose triples 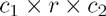 express the connection between a relation and its domain (*c*_1_) and range (*c*_2_) concepts. Objects 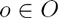 and links 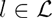 are instances of concepts 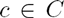 and relations 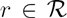, respectively. *Fig.* 3 shows an example of a phylogenetic processual ontology representing workflows abstract components.

**Figure 3.**
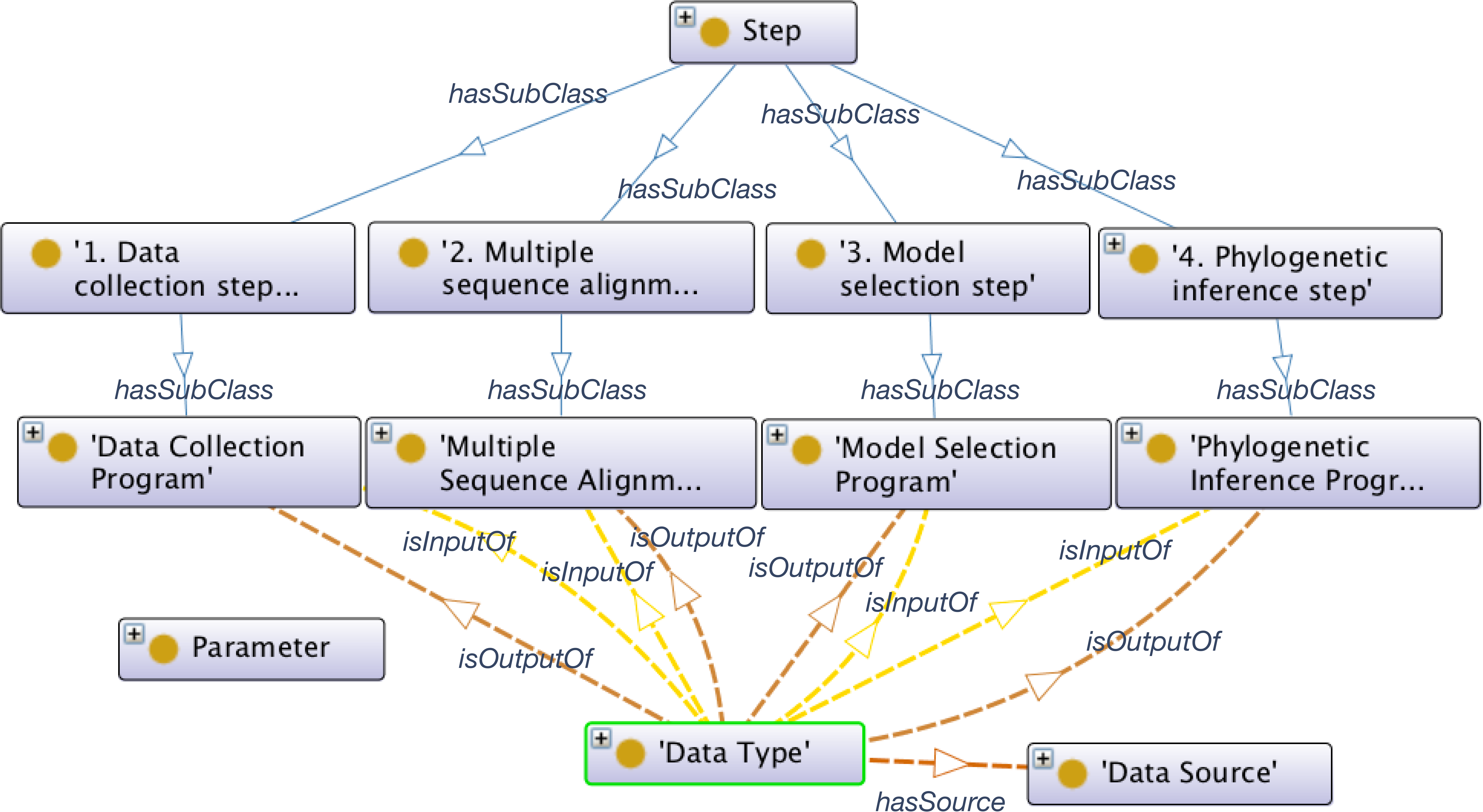
A sample from the proposed PHAGE ontology published in the BioPortal repository (see http://bioportal.bioontology.org/ontologies/PHAGE). Grey rectangles represent workflow reference concepts to: steps, programs, data types, resources and parameters. The hard arcs represent hierarchical *hasSubClass* relations and the dotted arcs represent input/output properties

#### 2.2 Workflow extraction from texts

Below, the sub-tasks of our workflow extraction solution are described. Following a standard Natural Language Processing (NLP) pipeline, we are going from morphological analysis of the text to the identification of grammatical classes and recognition of domain concept occurrences. The remainder addresses the known semantic ambiguity issues in a semi-automated way. Our approach, as an ontology-based annotation, is about learning how to recognize and classify workflow components (data and control flows) in texts using the ontological categories.

##### 2.2.1 Morphosyntactic annotation

The initial step divides the text into lexical items called tokens. In fact, the tokenisation operates words with white spaces between them. In our context, due to the specificity of the grammar used in bioinformatics texts, a specialized biomedical tokenizer is used relying on a dedicated pattern for bioinformatics tokens [30]. The text is also segmented into sentences by recognizing a set of known punctuation patterns (given as *regex*). The extracted tokens are further tagged with their appropriate grammatical classes, *a.k.a.* part-of-speech (POS) taggings. To this end, we use the MedPost POS tagger [31] which is based on a HMM (Hidden Markov Model) model trained over several biomedical MEDLINE texts. This tagger was able to achieve high accuracy (97.43% on 26 566 tokens) by using the contextual information in the HMM to resolve grammatical ambiguities.

##### 2.2.2 Semantic annotation

A specific OWL ontology gazetteer (terminology) is used in order to recognize bioinformatics concept and property instances in the text. This gazetteer was generated from widely-used databases: Gene Ontology [32], UniProtKB [33], NCBI Genbank [34] and the Joe Felsenstein Web site ^[5]^. Gene Ontology, NBCI Genbank and UniProtKB offer structured and up-to-date databases widely popular in the context of bioinformatics. We extracted scientific terms and their synonyms (common names, short/long forms, *etc*.) using ETL (Extract, Transform and Load) tools [35]. Felsenstein’s Web site offers also a very interesting categorization of 392 phylogeny packages and 54 free web servers where each package and web server lists a set of services and programs. Entries in this site are frequently (~6 months) updated. We crawled HTML contents from this web site using simple XPATH extraction rules in the Rapidminer platform [36]. Generated terms from the different databases and HTML pages are first stored in a relational database as a common storage before transforming them into an ontology using the schema provided by an expert (see *Fig.*3) and the SQL-SPARQL mapping tool ONTOP [37]. Also, we used the Sesame ontology repository ^[6]^ as a knowledge base manager. The resulted terminology is used to recognize term tokens and N-grams in texts and support the reconstruction of workflow sequences. This process faces known hurdles rooted in the semantic ambiguities that we address in the following paragraph.

##### 2.2.3 Disambiguation rules

For this purpose, we choose to create manually specific pattern rules to concepts grounding special semantic ambiguities *i.e.* polysemy [38]. For example, the term ‘MEGA’ is a polysemic term in our context: it could be either the common term (huge) or the name of a general purpose program. ‘MEGA’ could be used in sequence alignment, phylogenetic tree inference or/and in other phylogenetic tasks. To solve this type of ambiguities, we first created a set of disambiguation rules in JAPE ^[7]^. These rules enabled the construction of a large set of terms already annotated (classified) to be further integrated in a machine learner (see next section). For example, to verify if ‘MEGA’ was used in the inference step, we search for the verb ‘infer’ in the context of ‘MEGA’. By context here we mean the sentence in which the term appears (see Fig. 4). However, a sentence could carry a lot of information and has different forms (active or passive). Thus, we use a syntactic analyzer to construct a 2-levels parsing tree (chunk) [39] to detect the appropriate context of the term ‘MEGA’ in order to fire JAPE rules. For example the rule M searches if the term ‘MEGA’ was used to infer phylogenetic trees. *Rule M* (*sentence*) : { if the verb’s root in ‘MEGA’ chunk is ‘infer’ or there is a phylogenetic approach in the object of the chunk then create ‘PhylogeneticInferenceProgram’ Annotation}. The chunk of ‘MEGA’ is the verb phrase in the sentence where ‘MEGA’ is its Object/Subject (depending on the active or the passive voice of the sentence). The sentence in *Fig*. 4 shows that ‘MEGA’ is used a phylogenetic inference program since it is preceded by the approach ‘MaximumParsimony’ in its context. For each polysemic ‘concept step’ (see Fig. 3), we create *n* disambiguation rules (where *n* is number of ‘concept steps’ in the ontology) in order to clarify the ambiguities of program classes and generate a training set to the final disambiguation model.

**Figure 4.**
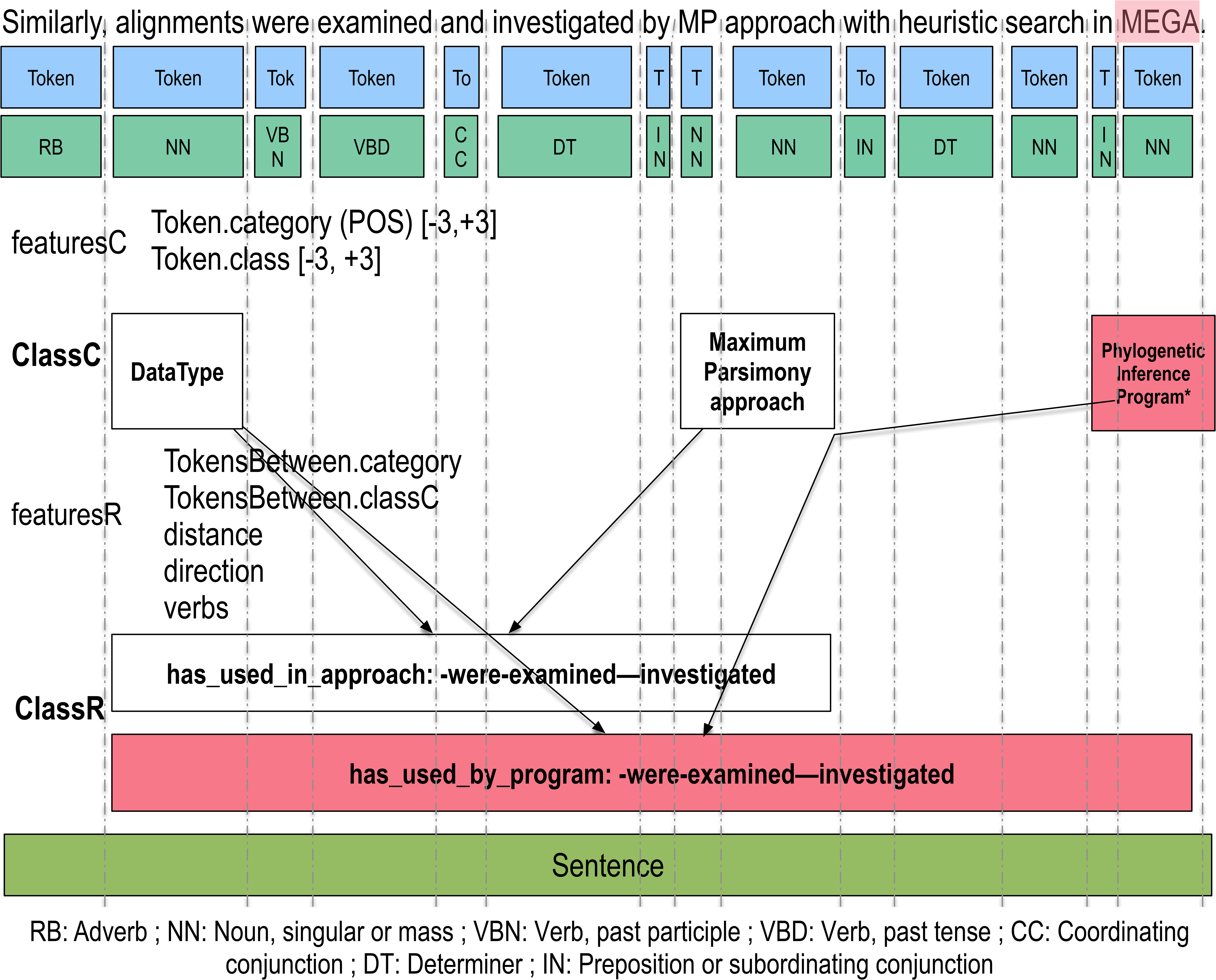
An example of concept and relation annotation in texts. After segmenting the text into tokens and POS tags, we construct a set of concept features *featuresC* to learn how to classify concepts *ClassC* and relation features *featuresR* to learn how to classify relations *ClassR*. Since our feature extraction method is context-based, we recognize each concept and relation in its sentence

##### 2.2.4 Learning features

A supervised learning approach is applied to ambiguity resolution. To this end, we created a set of features to support proper classification of bioinformatics concepts and properties in texts. For example in Fig. 5, in order to classify the term ‘MEGA’ (*ClassC*), we generate the set of features (featuresC) reflecting its context. The context covers a window made of: the *win*_*i*_ = 3 concepts preceding and the *win*_*i*_ ones following (*Token.class*) the concerning word (*e.g.* ‘MEGA’) in its context sentence. It also comprises the sequence of POS tags of the selected tokens (*Token.category*). In this example *Token.class* = {*DataType, MaximumParsimony Approach*} and *Token.category* = {*Determiner, Noun, preposition*}.

**Figure 5.**
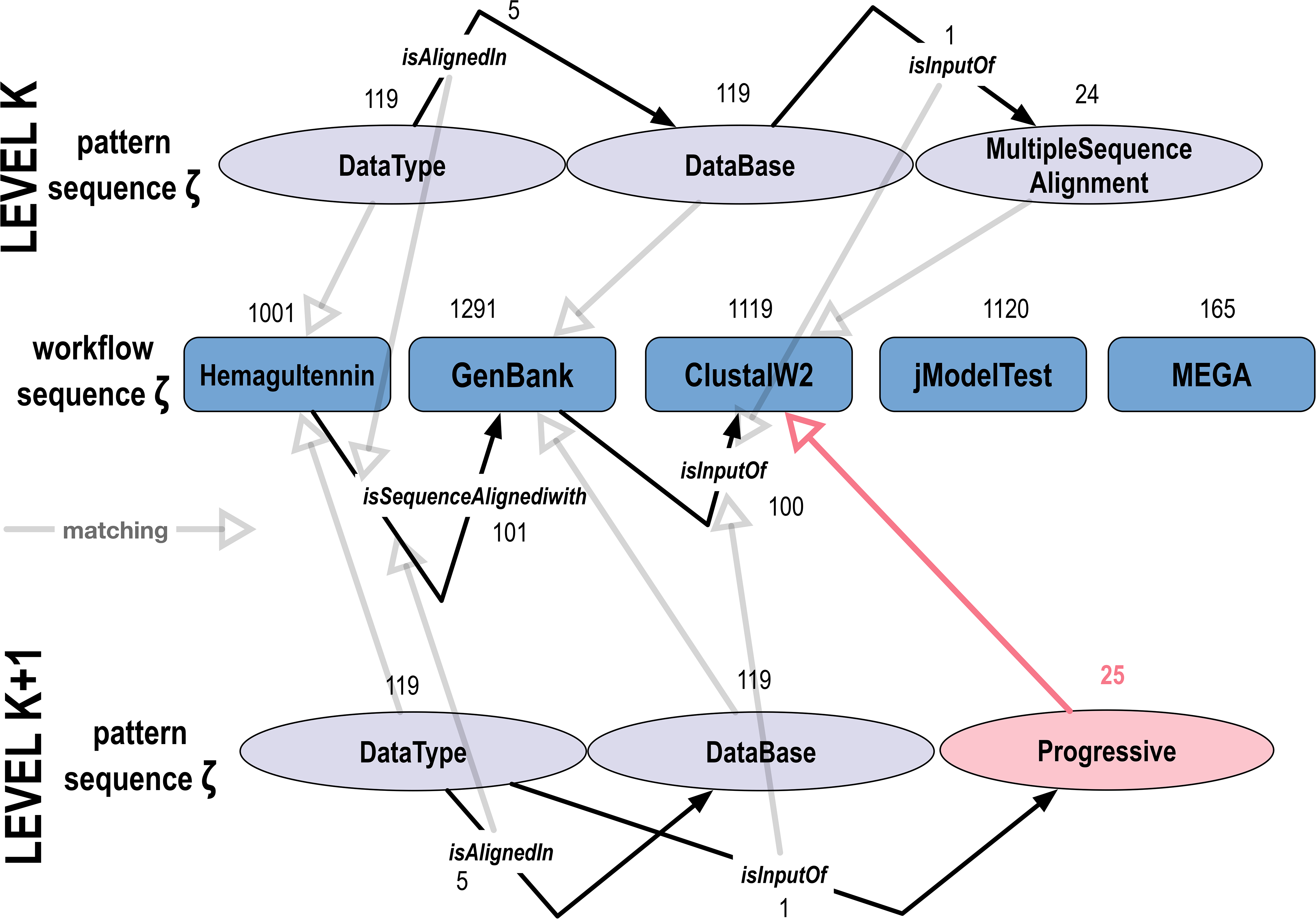
An example of pattern refinement from a level *k* to *k* + 1. Let *p* be an abstract workflow pattern (grey ovals) and *w* the concrete workflow sequence (blue rectangles). After refining/replacing the concept 24: *MultipleSequenceAlignment* with its immediate ontological child concept 25: *Progressive*, we need to verify if the matching between the concepts and relations of *p* and *w* still hold from the level *k* to *k* + 1

In addition, in order to classify a relation between the software ‘MEGA’ (domain) and the data ‘alignments’ (range) - *classR,* we define its context as the set of tokens between the domain and the range. Now, features (*featuresR*) represent token POS tags (*TokensBetween.category*), distance between domain and range, relation direction and the list of verbs between the concerned terms. The overall set of features is then used in the construction of a statistical model for automated annotation of the texts. Relation types between terms are already defined in the schema of the ontology (see Fig. 3). Tuples of concepts are then queried in order to recognize properties in text and calculate its features. For example, the ontological property ‘has_used_by.program’ between datatypes and phylogenetic inference programs is used to calculate the set of features *featuresC* representing the relation’s context in the sentence.

##### 2.2.5 Model creation and application

In this study, we choose PAUM as a classifier over others *i.e.:* CRF (Conditional Random Fields), SVM (Support Vector machines) and Random Forest. We compared PAUM models with the mentioned algorithms in terms of f-measure and speed, but we only present here the best classifier results. The PAUM algorithm (Perceptron Algorithm with Uneven Margins) [40] was used to learn term and relation classifications from the extracted features. PAUM was designed especially for imbalanced data and has been successfully applied to various named entity recognition problems: [41, 42]. PAUM differs from other classifiers as it approximates the best distance between positive and negative sets. As our approach is based on concept contexts to disambiguate polysemic words, the distance between surrounded concept examples (negatives) and the concerned word is important, same for domain and range relations. PAUM uses a specific margin function to adjust weights of positive and negative example features and thus shows competitive results. This function updates weights *w* for each example *x*_*i*_ based on a fixed bias parameters *τ*_*+*_ and *τ*_ representing the number of positive and negative examples, respectively. After creating and tuning our models to annotate the texts automatically, we applied a 10 folds cross validation analysis [43]. Extracted terms and links are then assessed using classic evaluation metrics *i.e.:* precision, recall and f-measure.

##### 2.2.6 Workflow reconstruction

From recognized links between concept instances in text, we build our concrete workflow knowledge database where items are the named entities ordered by the top-level pattern in the domain ontology. Concept and property instances serve to identify respectively the data-flow and control-flow of workflows. For example, from the tuples: (*ρ*_1_) (alignment, “has_dataSource”, hemagglutinin), (*ρ*_2_) (alignment, “used_in_SequenceAlignmentProgram”, ClustalW) and (*ρ*_3_) (alignment, “used_in_PhylogeneticInfence Program”, MEGA), we construct the workflow sequence S1 : 〈hemagglutinin, alignment, ClustalW, MEGA〉. We flatten then the workflow structure into a sequence of data and programs enriched with semantic relations (tuples) in order to represent the data and control flows.

##### 2.2.7 Workflow quality criterion

In order to evaluate the extracted concrete workflows, we calculate the similarity measure 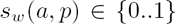 which represents the similarity between the actual *a* and the predicted *p* workflow components. If 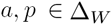 and *w* = (ζ, Θ), where *ζ* is the sequence of workflow items *i* at each step *j*, Θ is the set of triples 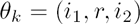 connecting relations *r* to the object domain at position *i*_1_ and the object range at *i*_2_, then the similarity *s*_*w*_ between *a* and *p* is given by the following formulas:

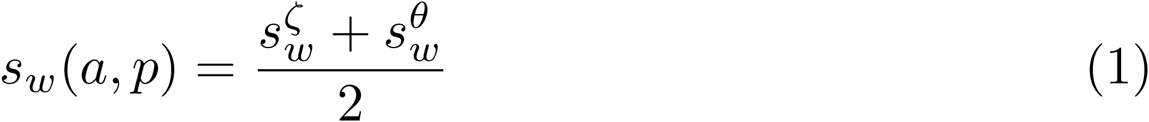

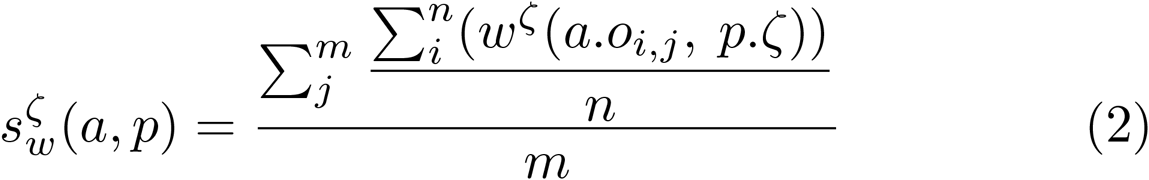

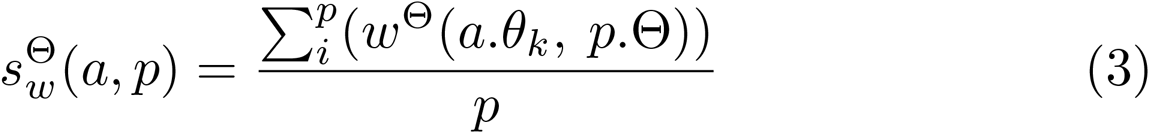

where *m, n* and *p* are, respectively, the total number of steps, items per step and relations in a workflow. Each object *o*_*i,j*_ is weighted by the accuracy measure *w*^*ζ*^ and each property *θ*_*k*_ is weighted by the accuracy 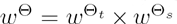 where the weight 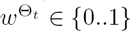 is the degree of a concept term accuracy and 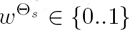 is the degree of a relation term accuracy. The similarity ratio of a workflow is then the mean of its components accuracies. Table 1 shows the weights that we propose to measure the degrees of accuracies of workflow components. If a term (concept or relation) in the actual workflow *a* is well annotated, then, its weight *w* is 1. Otherwise, if a term is partially correct, we assign a weight of 0.6. In the case where a relation type is partially correct, a much lesser weight is given (0.1). Finally if a term/step is not correct or absent, the weight is 0. These weights are inspired from the work of [43] where lenient precisions and recalls are calculated given a partial overlap measure.

**Table 1.** Objects and relations weights

| Weight | Description | Val |
| --- | --- | --- |
| $w_1^{\zeta_t}$ | The object term $o_{i,j}$ in $p.\zeta$ is correct. | 1 |
| $w_2^{\zeta_t}$ | The object term $o_{i,j}$ in $p.\zeta$ is partially correct. | 0.6 |
| $w_3^{\zeta_t}$ | The object term $o_{i,j}$ in $p.\zeta$ is not correct. | 0 |
| $w_1^{\Theta_t}$ | The relation term of $a.\theta_k$ in $p.\Theta$ is correct. | 1 |
| $w_2^{\Theta_t}$ | The relation term of $a.\theta_k$ in $p.\Theta$ is partially correct. | 0.6 |
| $w_3^{\Theta_t}$ | The relation term of $a.\theta_k$ in $p.\Theta$ is not correct. | 0.1 |
| $w_1^{\Theta_s}$ | The domain and range terms of $a.\theta_k$ in $p.\Theta$ are correct. | 1 |
| $w_2^{\Theta_s}$ | The domain or range terms of $a.\theta_k$ in $p.\Theta$ are correct. | 0.6 |
| $w_3^{\Theta_s}$ | The domain and range terms of $a.\theta_k$ in $p.\Theta$ are not correct. | 0 |

Weights values represent strings/substrings matchings between predicted item and triple terms against the gold standard terms.

#### 2.3 Generalized pattern mining

Next, we generate more abstract workflows from the set of concrete workflows extracted from the texts, i.e. the workflow database 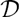.

##### 2.3.1 Candidate generation

Two descriptive languages are defined to represent both concrete workflows and abstract workflows. A data language ∆ is derived from workflow sequences *W* translated into IDs from the proposed ontology Ω. The second language Γ is used to represent patterns *p* in a canonic language [29]. Γ_*W*_ comprises generalized DAG patterns *p =* (ζ, Θ) where *p.ζ* is the sequence of concepts and *p.*Θ is the set of relation triples.

Our Apriori-like [26] level-wise mining approach considers the global pattern space as being made of levels. A level is defined *w.r.t* the overall depth of the pattern components within the ontology. Pattern candidates of a level *k* + 1 are generated from the *k*^*th*^-level frequent pattern by a refinement operator [44]. Here we extend the work of Adda et al. [29] by applying five different canonic operations to refine a pattern *p:* 1) append a root concept to a new transaction (step) in *p.ζ,* 2) add a root concept *to p.ζ,* 3) replace a concept from *p.ζ* by a direct instantiation (specialization) thereof, 4) add a root property between two concepts in *p.θ* and 5) replace a property from *p.θ* by a direct instantiation. For example (see Fig. 5), a possible specialization of the concept 24-’ *MultipleSequenceAlignmentProgram*’ in a pattern *p,* would be to replace it by one of its immediate successors in the ontology, say 25-’*Progressive*’.

The starting point is the computation of frequent level 1 patterns directly from the ontology (made of ontological root concept singletons). For each of the subsequent levels *k*+1, the level candidates are generated from frequent patterns at level *k* by applying canonic operations to them. The resulting candidates are first tested for the known infrequent sub-patterns and then the surviving candidates are sent to the database 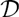 for a direct evaluation of their interestingness.

##### 2.3.2 Pattern quality criterion

As an approximation of the interestingness of a pattern, we use its support (frequency) *σ.* Our matching operation confronts a pattern *p*_*s*_ to a concrete workflow *w*_*s*_. If the matching holds, then *w*_*s*_ is in the support set of *p* in 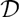, so it can be counted in the numeric value of the support criterion. The precedence relation, in candidate generation order which is based on instantiation, is considered in the matching data structure so we don’t need to recalculate the already matched sequences in each iteration. The matching verification begins from the last canonic operation on a pattern candidate. The matching data structure stores all previous matching IDs of a workflow pattern, searches and verifies whether the last refinement operation preserves the matching status or not. At a post-processing step, and in order to facilitate the interpretation by experts, concept and property IDs are replaced in each workflow pattern by the corresponding names from the ontology (labels).

Besides the support (frequency) as an interestingness feature, we calculate 6 other qualitative features. To characterize generalized workflow patterns, we propose to use representational features so we can measure the generalization/specialisation power of a pattern and its coverage. Hence, we propose to use the following set of features.

- **Specialization score** (*s*_*s*_). We measure here the generalization / specialization tradeoff score of a pattern. This score calculates the sum of its components levels (maximum depth of a concept/relation) from the ontology. Hence, a specialization score of 1 means that a pattern is too general. This score is weighted by the *α* ∈ {0…1} parameter to specify the ontological tradeoff level in which the user considers as the threshold level of generality. *α =* 1 means that the minimum generality level expected by the user is the root level in the ontology.
- **Number of steps** (*n*_*s*_). This feature calculates the number of steps covered by a pattern.
- **Number of data** (*n*_*d*_). This feature calculates the number of data used in a pattern.
- **Number of metadata** (*n*_*m*_). This feature calculates the number of annotations over pattern components. A metadata is an additional information on data or programs (parameters) discovered in the ontology.
- **Number of programs** (*n*_*p*_). This feature calculates the number of program types used in a pattern.
- **Number of relations** (*n*_*r*_). This feature calculates the weight of the data flow in a pattern. It calculates the number of triples (|Θ|) in a pattern.

Finally, patterns are ranked using the ranking function *r*(*p*) defined in:

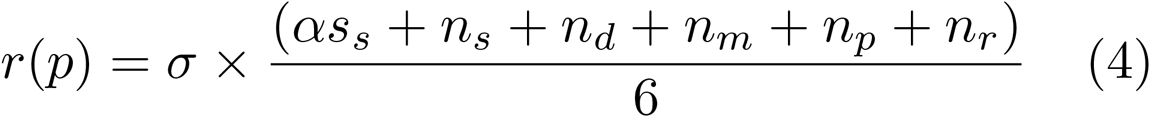

The ranking score measures how a pattern is informative in terms of number of information and generality it grounds. Note that *r*(*p*) is not normalized and if a workflow pattern has a high rank score, then it covers an important number of steps and additional information and is specific enough to be interpret by humans. The level of a pattern generality should be determined by the user using the *α* threshold level. In our experiments, we fixed *α* to 1.

### 3 Results

#### 3.1 Dataset

The experiments are derived from a sample of 1000 recent articles, journal and conference papers having at least one section titled: *’phylogenetic analysis*’ or *’phylogenetic analyses’.* However, these sections may have not sufficient information to reconstruct workflows from texts. Thus, we filtered the downloaded articles based on the smallest pattern that any phylogenetic analysis should represent. A phylogenetic study might use at least one data type and a phylogenetic inference program (see Fig. 3). Finally we retained a set of 332 articles. All the papers are collected from the digital repository PubMed Central (PMC) in XHTML format using the following query: *”((”open access”[filter] AND ”2015/01/01”[PDat] : ”2015/04/18”[PDat]) AND ”Phylogenetic analysis”[Section Title]) OR ”Phylogenetic analyses”[Section Title].*

Our public PHAGE ontology on BioPortal is composed of 37 different object properties (relations), 47,031 concepts (with 4 root concepts and 111 unique concepts) and more than 1,000,000 unique individuals (instances) without counting synonyms. Root concepts represent data types, data sources, steps and parameters. Step concepts represent the seven steps of phylogenetic analysis and relations between concepts represent data flows and control flows. Data source concepts are provenance metadata (database, GO, organisms, etc.) on workflow data components. We remind that this ontology has been collected in a semi-automated way from well-known databases such as NCBI, GeneOntology, UniProtKB and Felsenstein’s Web site.

#### 3.2 Extracted workflows from texts

A human expert has annotated manually 100 texts to construct a workflow test database. This set represents about 1/3 of the available set (the remaining 2/3 is used in the training process). The test set serves as a gold standard for the evaluation step. The expert annotates the given texts using our domain ontology schema on the platform GATE ^[8]^. The expert has annotated only terms of concepts and relations that serve to reconstruct phylogenetic workflows from texts. Next, we measure whether a term is annotated correctly, partially or unannotated at all (missing). We show in Table 2 details about the total counts of discovered annotations. We calculate the classic Information Retrieval f-measure [43] based on correct, partially correct and spurious annotations. In total, our system has annotated 2,561 concept terms while the expert has annotated 2,641. The overall average f-measure (average of strict and lenient f-measures) is 73%. We notice that 118 annotations (~ 20%) represent ambiguous program terms (GuessedProgram). In this case, the average f-measure is 84% with a precision of 73% and a recall of 98%.

**Table 2.**
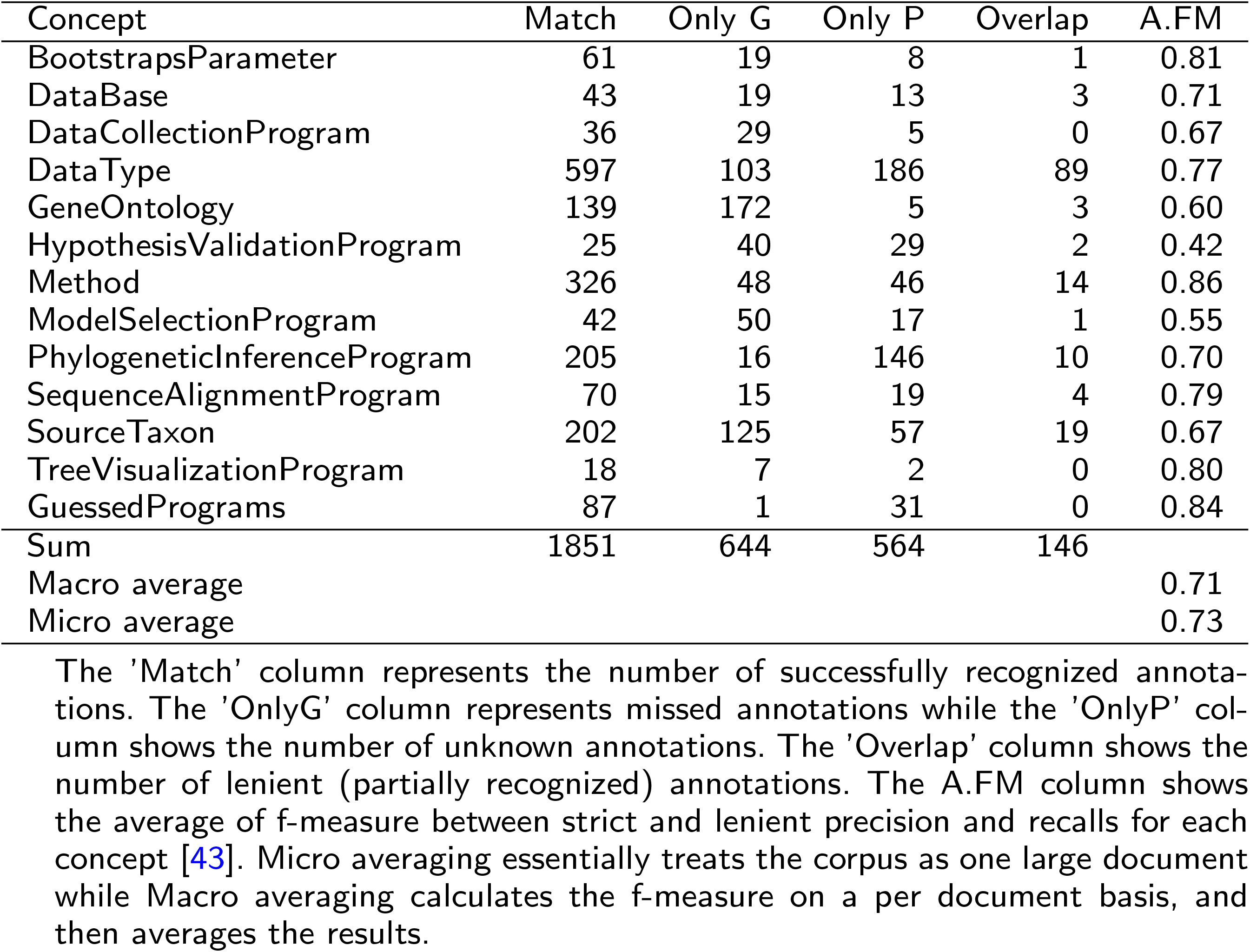
Concepts Evaluation Distribution

| Concept | Match | Only G | Only P | Overlap | A.FM |
| --- | --- | --- | --- | --- | --- |
| BootstrapsParameter | 61 | 19 | 8 | 1 | 0.81 |
| DataBase | 43 | 19 | 13 | 3 | 0.71 |
| DataCollectionProgram | 36 | 29 | 5 | 0 | 0.67 |
| Data Type | 597 | 103 | 186 | 89 | 0.77 |
| GeneOntology | 139 | 172 | 5 | 3 | 0.60 |
| HypothesisValidationProgram | 25 | 40 | 29 | 2 | 0.42 |
| Method | 326 | 48 | 46 | 14 | 0.86 |
| ModelSelectionProgram | 42 | 50 | 17 | 1 | 0.55 |
| PhylogeneticInferenceProgram | 205 | 16 | 146 | 10 | 0.70 |
| SequenceAlignmentProgram | 70 | 15 | 19 | 4 | 0.79 |
| SourceTaxon | 202 | 125 | 57 | 19 | 0.67 |
| TreeVisualizationProgram | 18 | 7 | 2 | 0 | 0.80 |
| GuessedPrograms | 87 | 1 | 31 | 0 | 0.84 |
| Sum | 1851 | 644 | 564 | 146 |  |
| Macro average |  |  |  |  | 0.71 |
| Micro average |  |  |  |  | 0.73 |

The 'Match' column represents the number of successfully recognized annotations. The 'OnlyG' column represents missed annotations while the 'OnlyP' column shows the number of unknown annotations. The 'Overlap' column shows the number of lenient (partially recognized) annotations. The A.FM column shows the average of f-measure between strict and lenient precision and recalls for each concept [43]. Micro averaging essentially treats the corpus as one large document while Macro averaging calculates the f-measure on a per document basis, and then averages the results.

We present in Fig. 6 the distribution of similarities *s*_*w*_(*a,p*) between the predicted and the gold standard concrete workflows. The overall similarity *s*_*w*_ is 0.83 over the 100 extracted concrete workflows. This figure shows also details about similarity distributions of both concepts and relations in workflows. The overall mean of relation similarity is 0.89 and concept similarity is 0.79. The best concept results are phylogenetic inference programs where the similarities are above 0.8 except for 5 workflows.

**Figure 6.**
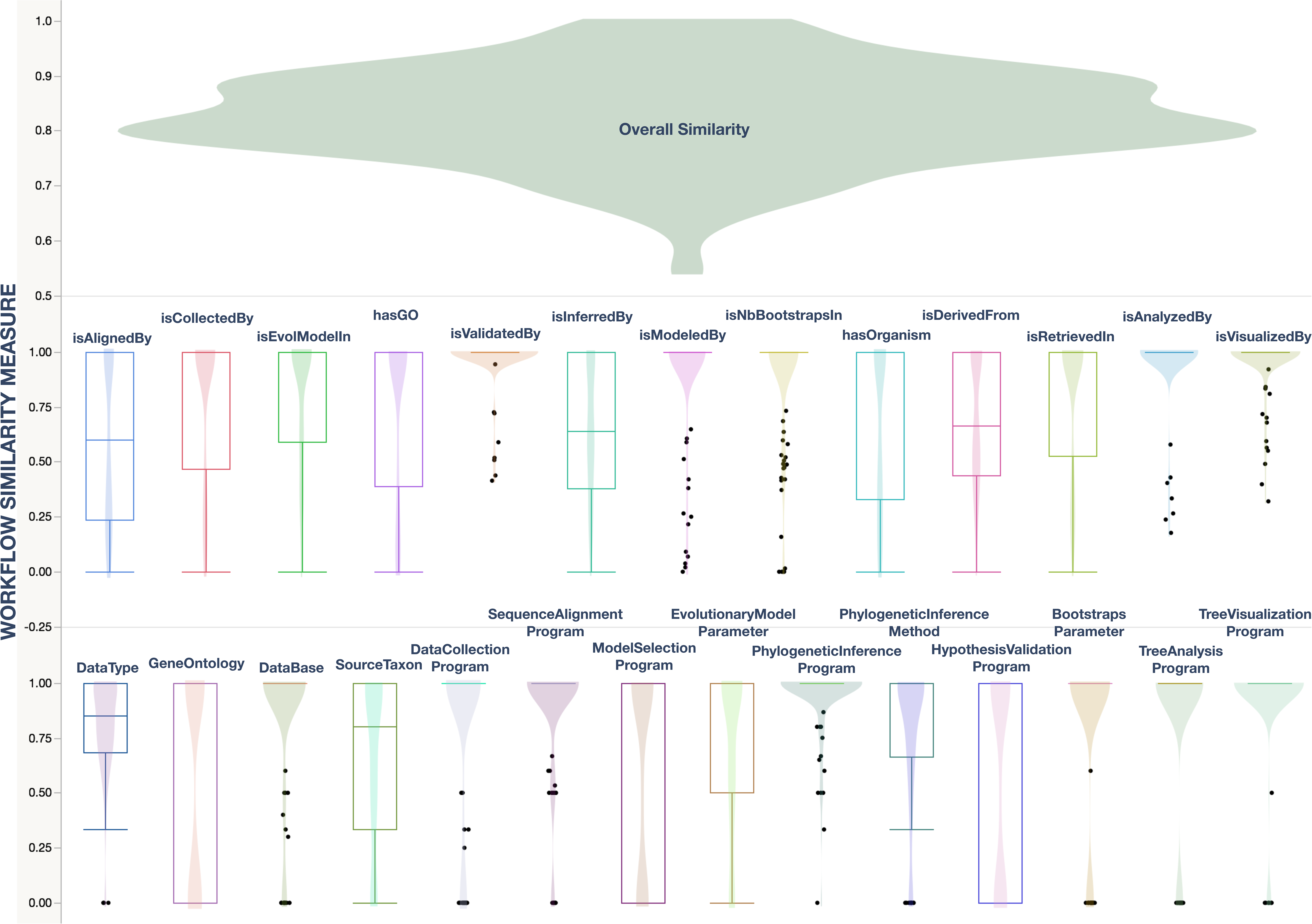
Workflow similarities results. The top figure represents the overall similarity between the actual *a* (annotated by the expert) workflows and the predicted *p* ones. The experiments have been performed on 100 recent articles downloaded from the PMC database. The second and third figures represent box-plots and contour distributions of concept and relation similarities between *a* and *p*. Bold points represent the interquartile ranges outliers

Moreover, in one side, gene ontology and taxa annotations in workflows are 0.6 similar within the actual workflows which is understandable since only full scientific terms are considered while tagging the name of species, genes and proteins in texts. Though, synonyms weren’t considered for the mentioned concepts. These are dismissed in order to minimize the error cost of the abstractions in common terms (e.g. cell, RNA, polymerase, etc.). In the other side, all program terms similarities are above 0.8 except for model selection and hypothesis validation programs (0.6). This is due to the context ambiguities within these categories. Bioinformaticians and biologists tend to use general purpose packages instead of specific tools for model selection and tree validation steps. The context of such tools are 40% of the cases out if its sentence area.

In the relation side, links between data types and tree visualization programs show the best similarities (the mean similarity is 1). Mean similarities of relations between : data types and MSA (Multiple Sequence Alignment) programs, data types and phylogenetic inference programs and data types and species are between 0.65 and 0.77. Both predicted relations between model selection programs and their model parameters, and predicted relations between bootstrapping programs and their number of replicates are 0.9 similar with the expected workflow relations.

#### 3.3 Generalized bioinformatics patterns

At the mining step, we use an Apriori-like sequential mining algorithm that exploits the domain ontology to output the most frequent generalized workflow patterns. In an unrestrained run, the algorithm scans the pattern space from the top downwards, at each step exploring the patterns at a specific generality level. After the first few levels, each step generates a considerable number of patterns. A well-known regularity with patterns over a hierarchy of items can easily be observed in our case: patterns of high frequency tend to be too general, i.e. involve concepts and properties from the higher levels of the ontology, and therefore have little practical value. Conversely, specific patterns -whose component concepts and properties lay close to the leafs of the ontological hierarchies- typically have low supports and thus might not make to the mining result. As an illustration, the diagram in Fig. 5 might be interpreted as follows: knowing that the subsequence: “database and multiple sequence alignment” is frequently used, doesn’t ensure that every local or global alignment program is frequent too. The question that we investigate next is about the optimal target level(s). If patterns specialized to this target are mined with concepts at the same ontological levels (homogeneous concepts), then we can skip the rest of the levels. Thus, we parameterized our algorithm to let bioinformaticians choose their preferred ontological level (*α).*

In the next experiments, we generated over 4 000 patterns using exclusively concepts from the 1^*st*^ level of the application ontology.

We evaluate the components of generated patterns using the qualitative features described in section 2.3. *Fig.* 7 shows results of the distribution of each feature on the set of generated patterns. The majority of patterns (89%) have high degrees of specialization (between 9 and 17). For low values of minimum of support (*σ* < 0.2), pattern levels are high, thus we obtain more specific patterns while decreasing *σ.* The distribution of the number of steps shows that our patterns cover mostly 6 steps of the phylogenetic inference analyses which is a very good coverage. In addition, these patterns have an average of 2 data per pattern and are annotated by up to 2 metadata. The number of programs follows the distribution of step frequencies and most of the discovered patterns have 6 ~ 8 relations between their components.

**Figure 7.**
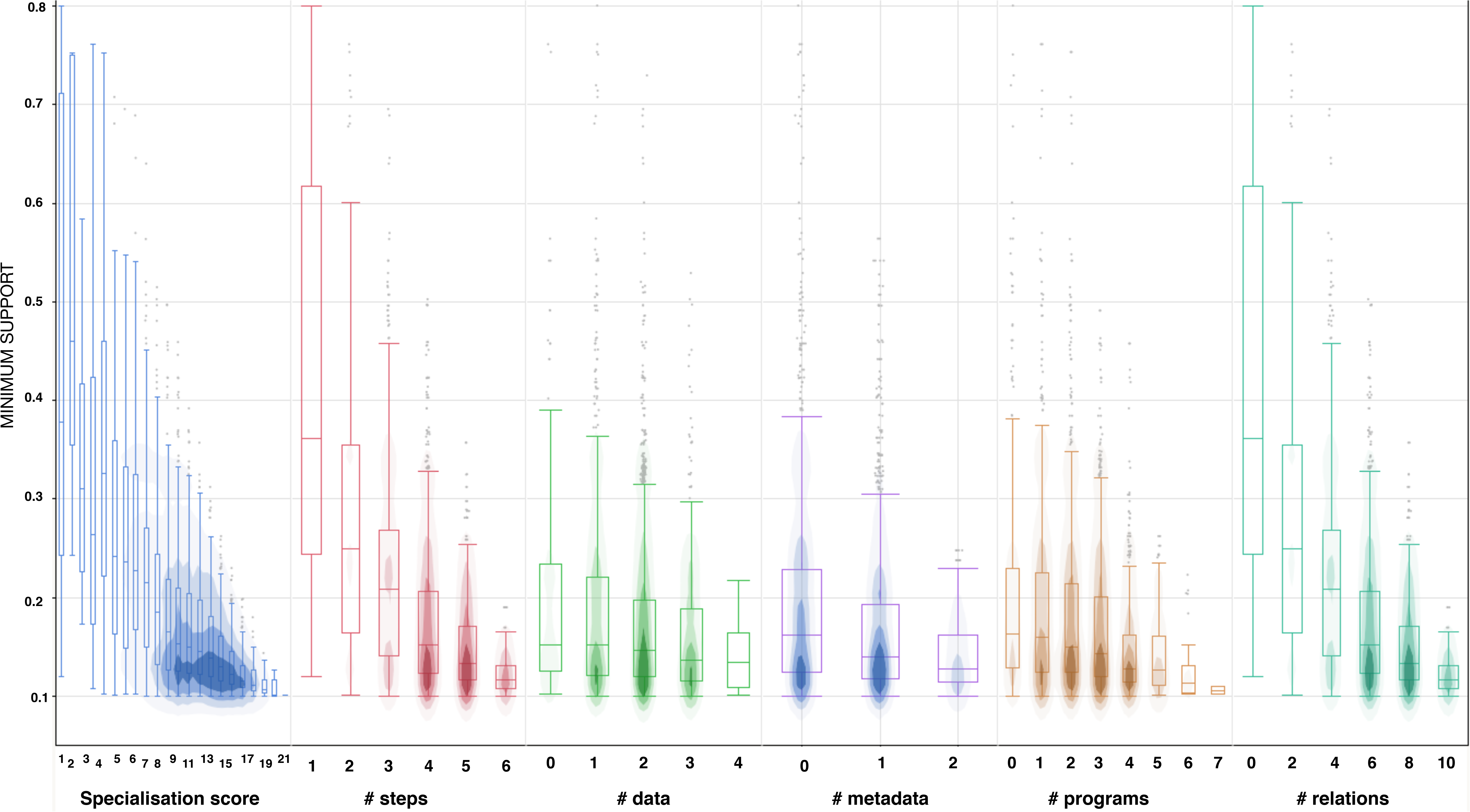
Pattern features distributions. This figure shows the frequencies of qualitative features over generated patterns using quantile box plots and contour densities. The Y axis shows the minimum of support used to generate patterns and the X axis shows feature values

Next, we evaluate generated patterns by measuring their ranking scores using the qualitative measures, so the more a pattern is informative (in terms of its support, degree of specialisation, number of covered steps, etc.), the highest rank it gets. Fig. 8 describes the percentage of ranking scores (rounded values) over the set of patterns. Top-40 best patterns (1%) have a rounded rank score of 10 (best score). This means that only 40 frequent patterns from the discovered ones are the most informative and cover the best values of features.

**Figure 8.**
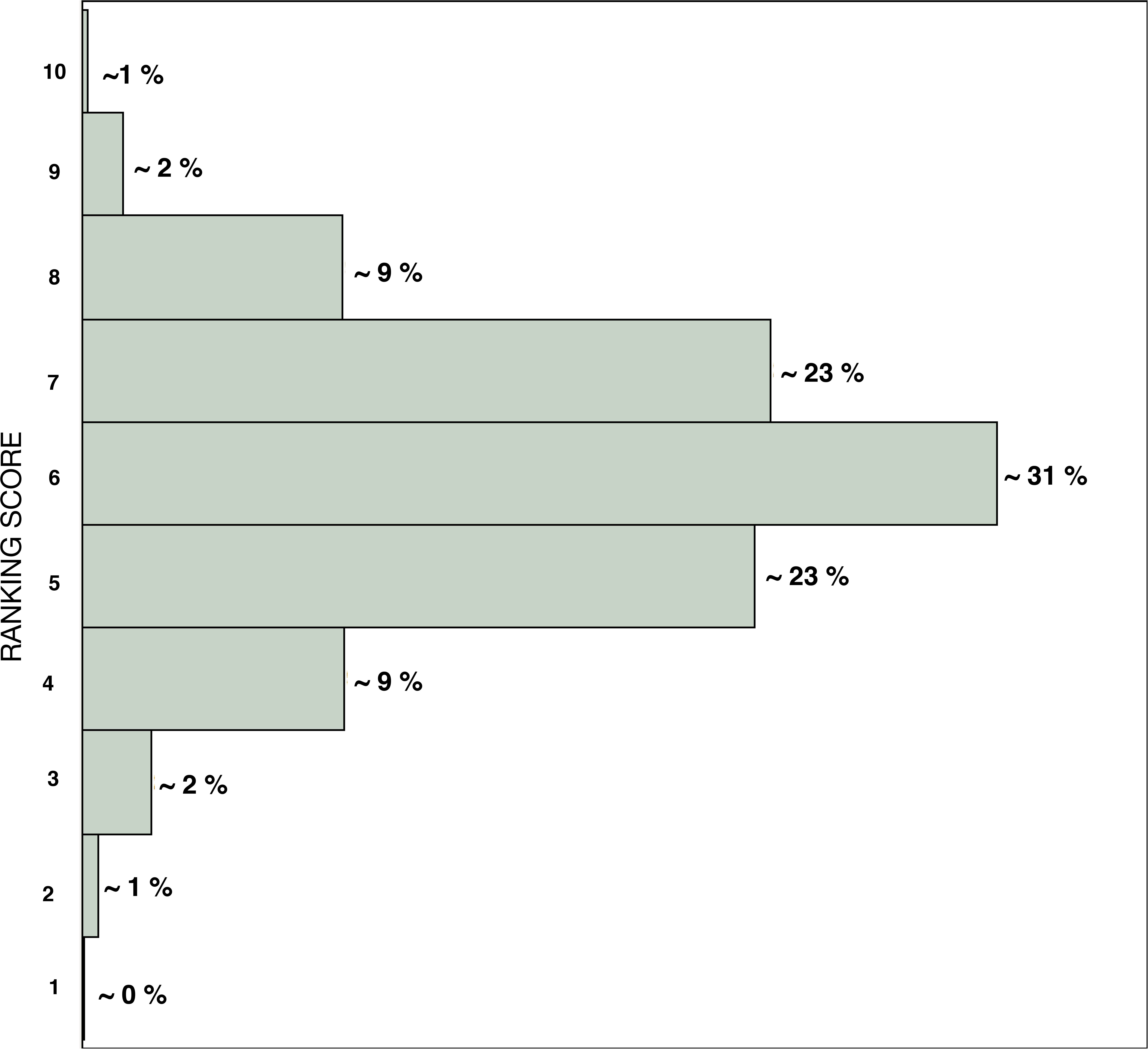
Pattern ranking score *r* distribution. This figure shows the distribution of *r* values over generated patterns in terms of percentages

We present, next, a sample of the discovered generalized workflows based on the concrete database.

P1 〈〈{Protein, Alignments, Tree, NCBI}, {Global MSA}, {NJ_P, NJ_M, CAT}, {Bootstrapping_P}〉, {(1, isRetrievedIn, 4), (2, isAlignedBy, 5), (3, isInferredBy, 6), 6, isDerivedFrom, 7), (8, isEvolModelIn, 6)}〉: [*σ =* 0.5]
P2 〈〈{Protein, Alignment, Tree, Plant, PDB}, { Gen-eral_MSA_P}, {ModelSelP, CAT}, {Bayesian_P}〉, {(1, hasOrganism, 4), (1, isRetrievedIn, 5), (2, isAlignedBy, 6), (1, isModeledBy, 7), (3, isInferredBy, 9), (8, isEvolModelIn, 7)}〉: [*σ =* 0.2]
P3 〈〈{Protein, Alignments, Trees, Virus, PDB}, { GlobalMSA_P}, {ModelSelection_P, JTT}, { Maximum_Likelihood_P}, {Bootstrap-ping_P}〉, {(4, isOrganism, 1), (1, isRetrievedIn, 5), (2, isAlignedBy, 6), (1, isModeledBy, 7), (3, isInferredBy, 9), (3, isValidatedBy, 10), (9, isEvolModelIn, 7)}〉: [*σ =* 0.1]

The first pattern *P*1 is 50% frequent in the concrete workflow database. Protein sequences are used with a global Multiple Sequence Alignment (MSA) programs and followed by a neighbour joining inference program and a bootstrapping program. These proteins are collected from the NCBI database and modelled with the CAT model. The second pattern is less frequent (20%) than the first one but it contains more information. Here *P*2 presents 4 phylogenetic steps. In the first step, plant protein sequences are downloaded from the PDB database. These sequences are then aligned using a General MSA program and modelled by the CAT under a model selection program. A Bayesian program is also used to reconstruct phylogenetic trees. The third pattern is quite frequent (10%) but it presents 5 different steps which is very interesting. Here we find that viral protein sequences are first loaded to be aligned with a global MSA program. Next, a model selection program is used to represent the evolutionary history of these sequences using the JTT model. The maximum likelihood approach is also used to infer the phylogenetic trees and a bootstrapping program to validate them.

## 4 Discussion

As shown in the previous section, our solution may produce truly interesting results, yet the traversal of the levels in the ontology structure and, to an even greater extent, the levels in the pattern space, may prove costly. Thus, if the appropriate levels in the ontology could be fixed beforehand, so that only the corresponding concepts and properties are used for pattern generation, the overall mining step will be performed much faster and the output is more likely to be of interest. The ultimate interestingness of a pattern depends on the decision-maker (subjective criteria) and does not solely depend on the statistical strength of the pattern (objective criteria, e.g. support) [45, 46]. Albeit useful, objective measures are not enough to determine the interestingness of a pattern particularly in complexly structured domains like phylogenetics problem solving. Our intended solution is to let users choose their levels of interest in the ontology. Then, the ranking score could be used to avoid checking too many patterns [46]. To the best of our knowledge existing solutions for semantic-based workflow mining [11, 15, 17, 18] propose domain-specific solutions while producing substantially different types of abstract models. Therefore, comparing our framework with systems like Proteus or eSysbio in terms of generated models, while conceivable, is not a trivial task. Furthermore, in order to validate the quality of our results, a human expert has already evaluated the building blocks of the generated workflows (see the above section 3). The abstract workflows produced here could be used in a pattern-based recommender to suggest a next tool to a user who got lost after a certain number of steps with the workflow manager platform. To that end, the system has to match the partial workflow corresponding to the user’s session against (the prefixes of) the abstract patterns from our pattern base. The remaining part (suffix) of a matching pattern represents a potential hint as to the way the user’s session might unfold from that step on. More specifically, if we take *Example 2.* as an illustrative example, we can apply the pattern *P*3 (see section 4.3) since it matches the sequence of the concrete workflow *S2* : 〈*rRNA_sequences, ClustalX, MAPPS*〉. *S*2 do matches with *P*3 in the data type ‘Protein sequences’ (same as in the source and database metadata) and in programs used: ‘Global MSA Program’ and ‘Model Selection Program’. From *P*3 we can recommend to use a JTT model, a maximum likelihood inference approach and a bootstrap-based tree validation for the next steps. With a concept-based recommendation, the system could work on different granularity levels and hence offer more flexible recommendations. Indeed, instead of objects as concrete tools and resources from concrete workflows, we suggest a class of objects to assist users with more semantically rich recommendations. Semantic suggestions could guide to compose more complex pipelines by presenting likely compositions. Users shouldn’t be worried about very specific details (software versions, technical supports, *etc*.) while constructing their workflows.

## 5 Conclusions

With many biologists and bioinformaticians having little experience of automating bioinformatics analyses, it is important to provide relevant recommendations in order to improve the efficiency of analyses. Concept-based pattern recommendation helps users not only to acquire contextual information but also to provide valuable suggestions. In this article, we presented the foundation of such recommendation system. We intend to extend our ontology-based mining framework in order to consider a variety of criteria, for example programs’ parameters (e.g. Gamma rates, transition ratios, number of bootstraps, etc.), more complex control structures (e.g. parallel executions, if-else statements) and resource quality in terms of time (the most recent tools/databases), author (focusing on domain’s experts), journals, popularity, etc,). Updating our text corpora with the most recent articles will help to produce more relevant patterns. Moreover, the proposed generalized patterns could be used to improve even journals quality to select the most relevant articles using the abstract and concrete workflow information within texts. For instance representing the proposed workflow used in a submitted article could also assist the reviewer in a semi automatic assessment and reusability of the proposed procedure.

## Declarations

### List of abbreviations

DAG: Direct Acyclic Graph ; DAML: DARPA Agent Markup Language ; GATE: General Architecture for Text Engineering ; KM: Knowledge Management ; NJ: Neighbor Joining ; NLP: Natural Language Processing ; MSA: Multiple Sequence Alignment ; OIL: Ontology Interchange Language ; OWLS-S: Web Ontology Language for Services ; PAUM: Perceptron Algorithm with Uneven Margins ; POS: Part-Of-Speech ; SM: Semantic Web ; WMS: Workflow Management System

### Ethics approval and consent to participate

Not applicable.

### Consent for publication

Not applicable.

### Availability of data and materials

Tools and data sources are available in: http://labo.bioinfo.uqam.ca/tgrowler

### Author’s contributions

AH implemented tools of workflow extraction and mining. PV and ABD examined the related work and contributed with AH in result analyses and interpretations. All authors read and approved the final manuscript.

#### Acknowledgements

Not applicable

## Footnotes

[1] Armadillo is a drag and drop workflow platform dedicated to phylogenetic as well as general bioinformatics analysis. www.bioinfo.uqam.ca/armadillo

[2] www.ebi.ac.uk/Tools/emboss

[3] www.esysbio.org

[4] www.bionerds.sourceforge.net

[5] www.evolution.genetics.washington.edu

[6] www.rdf4j.org

[7] Java Annotation Patterns Engine: www.gate.ac.uk/sale/tao/splitch8.html

[8] Gate is a cost-effective environment for annotation projects using NLP tools. www.gate.ac.uk

